# Characterization of a thermal stable poliovirus mutant reveals distinct mechanisms of virion stabilization

**DOI:** 10.1101/410175

**Authors:** Y Nguyen, Palmy R. Jesudhasan, Elizabeth R. Aguilera, Julie K. Pfeiffer

## Abstract

Enteric viruses, including poliovirus, are spread by the fecal-oral route. In order to persist and transmit to a new host, enteric virus particles must remain stable once they are in the environment. Environmental stressors such as heat and disinfectants can inactivate virus particles and prevent viral transmission. It has been previously demonstrated that bacteria or bacterial surface glycans can enhance poliovirus virion stability and limit inactivation from heat or bleach. While investigating the mechanisms underlying bacterial-enhanced virion thermal stability, we identified and characterized a poliovirus mutant with increased resistance to heat inactivation. This poliovirus mutant, M132V, harbors a single amino acid change in the VP1 capsid-coding that is sufficient to confer heat resistance. M132V was comparable to wild-type virus for replication *in vitro*, as well as for replication and pathogenesis in orally-inoculated mice. Although the M132V virus was stable in the absence of bacteria or feces at most temperatures, M132V virus was stabilized by feces at very high temperatures. Additionally, the M132V virus does not have enhanced stability during bleach treatment, suggesting that thermal and bleach inactivation mechanisms are separable. Our results suggest that different mechanisms underlie virion stabilization by bacteria and the M132V mutation. Overall, this work sheds light on factors that influence virion stability.

**IMPORTANCE:** Viruses spread by the fecal-oral route need to maintain viability in the environment to ensure transmission. Previous work indicated that bacteria and bacterial surface polysaccharides can stabilize viral particles and enhance transmission. To explore factors that influence viral particle stability, we isolated a mutant poliovirus that is heat resistant. This mutant virus does not require feces for stability at most temperatures, but can be stabilized by feces at very high temperatures. Even though the mutant virus is heat resistant, it is susceptible to inactivation by treatment with bleach. This work provides insight into how viral particles maintain infectivity in the environment.

## INTRODUCTION

Enteric viruses such as poliovirus (PV) cause mild to severe diseases in humans. PV is a non-enveloped, single-stranded 7.5-kb positive-sense RNA virus in the *Enterovirus* genus of the *Picornaviridae family*. Spread by fecal-oral route, PV replicates in the gastrointestinal tract and can disseminate and cause neuronal damage and subsequent paralysis in the host. Although highly successful vaccines prevent poliomyelitis in most countries, PV serves as a useful model system to understand fundamental aspects of virology.

PV’s 30-nm icosahedral capsid is comprised of 60 copies of each of the viral capsid proteins VP1, VP2, VP3, and VP4 (1). VP1, VP2, and VP3 form a network that encompass the surface of the capsid while VP4 is on the interior of the virion (1, 2). VP1 contains a highly conserved lipid moiety pocket that is associated with virion uncoating (3). Like many non-enveloped viruses, PV’s structure is dynamic at physiological temperatures, and internal capsid amino acids are reversibly exposed in a process called “breathing” (4-6). These transient events may be an important precursor to the normal entry process. However, these conformational changes can also lead to irreversible premature RNA release prior to cell entry.

Previous work elucidating capsid stability determinants has led to the identification of thermal stability mutants and stabilizing compounds (7-11). For example, Adeyemi *et al.* identified two capsid mutations in VP1 (V87A and I194V) that enhance virion stability by limiting premature uncoating and release of viral RNA at high temperatures (9) and work from Shiomi et al. also suggested that VP1 mutation V87A contributes to virion heat resistance (8).

Because PV is transmitted by the fecal-oral route, PV particle stability in the environment is crucial for viral persistence and transmission to new hosts. Furthermore, recent studies revealed that the intestinal microbiota promote PV infection by increasing virion stability and cell attachment (12-14). A PV mutant with reduced lipopolysaccharide binding (VP1-T99K) had a transmission defect, exhibiting reduced stability in feces compared to wild-type PV, without displaying any defects in cell attachment and replication *in vitro*, or pathogenesis in mice. Moreover, it was determined that PV interacts with bacteria by binding to bacterial surface glycans including lipopolysaccharide and peptidoglycan (12, 13), enhancing virion stability by preventing premature RNA release (12). While it is clear that binding to bacteria or bacterial surface glycans enhances thermal stability of PV virions, the specific interaction determinants between bacterial components and virion capsids—and therefore the mechanism of thermal stabilization by bacteria--remain unknown.

Here, we exploit a PV mutant that is thermostable in the absence of bacteria to uncover mechanisms of bacterial-mediated PV thermostability. By selecting for variants that maintained infectivity after heat exposure, we identified a single amino acid substitution in the VP1 viral capsid protein, VP1-M132V, that is sufficient to increase virion thermal stability, yet does not alter PV replication, shedding, or pathogenesis in mice. We found that although the M132V mutant is generally stable independently of bacteria or feces, at higher temperatures fecal contents rescued M132V viruses from thermal inactivation. Finally, we revealed that the M132V mutation did not protect the virion from other environmental stressors, such as bleach inactivation, suggesting that the M132V phenotype is specific to heat resistance

## RESULTS

### Identification of a thermostable PV mutant

Previously we found that binding to bacteria or bacterial surface polysaccharides can reduce thermal inactivation of PV (13) (12). Additionally, we showed that a single amino acid substitution in the VP1 viral capsid protein, T99K, diminished lipopolysaccharide binding and virion stabilization by lipopolysaccharide (12). To further define factors that influence virion stability, we sought to select for PV mutants with increased thermal stability in the absence of bacteria by using repeated exposure to elevated temperatures followed by amplification of viable viruses. PV was incubated at 43°C in PBS for 6 h followed by quantification of viable virus by plaque assay. Under these conditions, the infectivity of WT PV was reduced by 99.99% compared to no heat treatment controls (Figure 1A). Remaining viable viruses were amplified in HeLa cells followed by incubation at 43°C in PBS for 6 h. This cycle was repeated a total of 10 times. By passage 5 (P5), the heat exposed viruses had 2-fold increased viability after heat treatment compared to the WT virus, and by passage 8 (P8), the heat-exposed viruses had 500-fold increased viability after heat treatment compared to the WT virus (Figure 1A). After passage 10, we isolated RNA from the heat-passaged viruses, performed reverse-transcriptase (RT)-PCR, and sequenced the capsid-coding region. Sequence alignment revealed a single amino acid change, M132V in the VP1 capsid protein (Figure 1B). The VP1-M132V mutation is buried inside the capsid within the 8-stranded β-barrels of VP1 (Figure 1C).

**FIG 1.**
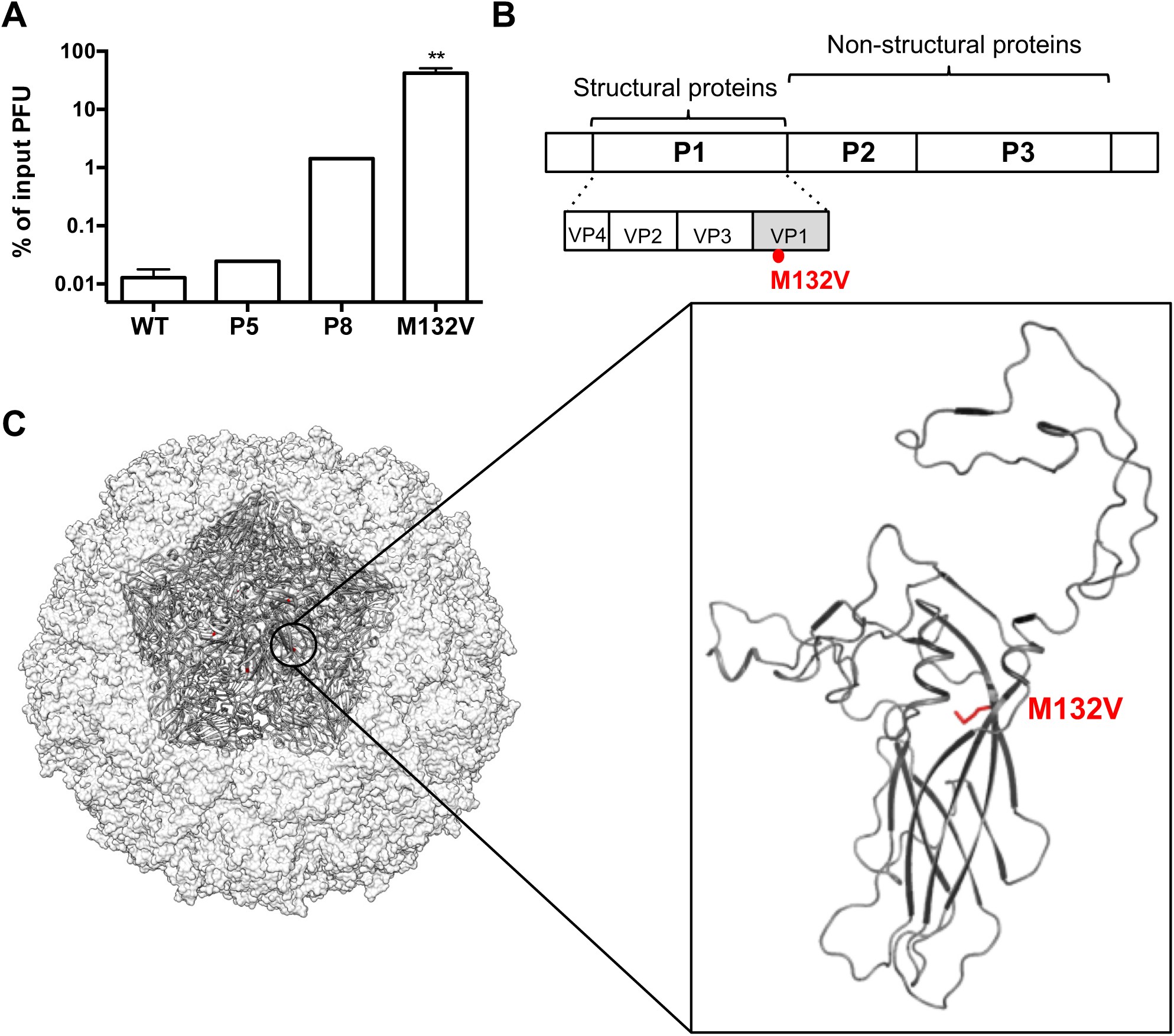
Selection and identification of a thermostable PV mutant. (A) Thermal stability profiles for WT PV, passage 5 and 8 viruses, and the VP1-M132V mutant. Viruses were incubated in PBS at 43°C for 6 h followed by plaque assays. Titers from time point zero were compared to post-incubation titers to determine the % input PFU. For WT and M132V mutant n=6; for the P5 and P8 viruses n=1 because the remainder of the samples were used for the next passage. (B) Genome schematic of PV with M132V mutation in VP1 indicated in red. (C) Poliovirus structure showing one 5-fold symmetry axis with the location of VP1-M132V highlighted in red. The insert shows a ribbon model of VP1 with the position of M132 amino acid shown in red. Data are mean ± SEM; **p<0.01.

### A single amino acid change, VP1-M132V, is sufficient for thermal resistance

To confirm that the VP1-M132V mutation confers thermal resistance, we cloned the M132V mutation into the PV infectious clone and examined viability by plaque assay after incubation at 43°C for 6 h. As shown in Figure 1A, the M132V amino acid change was sufficient to increase infectivity by more than 1000-fold compared to WT PV. As a second method to quantify virion stability using a physical/cell-free approach, we examined whether the M132V mutation alters viral RNA release using a Particle Stability Thermal Release assay (PaSTRY) (15) (12). Gradient-purified WT or M132V PV was mixed with SYBR Green and heated in a real-time PCR machine with fluorescence monitoring. SYBR Green binds viral RNA upon release from capsids and can be used to determine the temperature at which RNA release occurs. WT PV generated a peak of fluorescence intensity indicating RNA release at 51°C (Figure 2). However, for M132V PV the fluorescence intensity peak shifted to 78°C (Figure 2). Overall, these data indicate that the M132V mutation in VP1 enhances viral stability by limiting RNA release.

### Replication of M132V and WT PV is equivalent in cultured cells and in mice

Because the M132V mutation can increase viral stability by delaying RNA release during heat exposure and RNA release is critical for infection, we wanted to determine if this mutant has a growth defect compared to WT PV. We first compared the replication of M132V and WT PV in cell culture using single-cycle growth curve assays. HeLa cells were infected at a multiplicity of infection (MOI) of 10, and viral titers were determined over time by plaque assay. As shown in Figure 3A, the M132V mutation did not significantly affect viral yields in HeLa cells. Next we wanted to determine whether the M132V mutant has a fitness defect in mice by examining viral shedding and pathogenesis in orally-inoculated C57BL/6 PVR-IFNAR^-/-^ mice. These mice express the human poliovirus receptor (PVR) and are deficient for the interferon (IFN) α/β receptor (16), which confers oral susceptibility to PV. Mice were orally inoculated with 10^8^ PFU of either WT or M132V PV, fecal samples were collected at 24, 48, and 72 hours post-inoculation (hpi), and titers were determined by plaque assay. As shown in Figure 3B, we found that fecal titers were similar for WT and M132V viruses, although M132V titers were slightly lower than WT at 48 hpi. However, viral pathogenesis was equivalent for WT and M132V PV (Figure 3C). Taken together, these results suggest that the M132V mutation does not confer a major replication defect *in vitro* or *in vivo*.

### Feces can stabilize M132V PV at higher temperatures

Since PV spread by the fecal-oral route, virion stability in feces is important for transmission to a new host. Previously we demonstrated that feces stabilize WT PV (12). Because M132V PV has enhanced thermal stability, we suspected that M132V virions do not require feces for stability in the environment. To examine environmental stability of M132V and WT PV, we collected feces from uninfected C57BL/6 PVR-IFNAR^-/-^ mice, resuspended them in PBS, and incubated this mixture with 10^5^ PFU of M132V or WT PV at 37°C prior to quantification of viable virus by plaque assay over several time points. As shown in Figure 4A, WT PV was stabilized by feces over time, consistent with our previous findings that bacteria or bacterial components in feces enhance virion stability (12, 13). In contrast, M132V PV was stable even in the absence of feces, although fecal components slightly increased virion stability of the mutant at a late time point (day 8)(Figure 4A).

**FIG 2.**
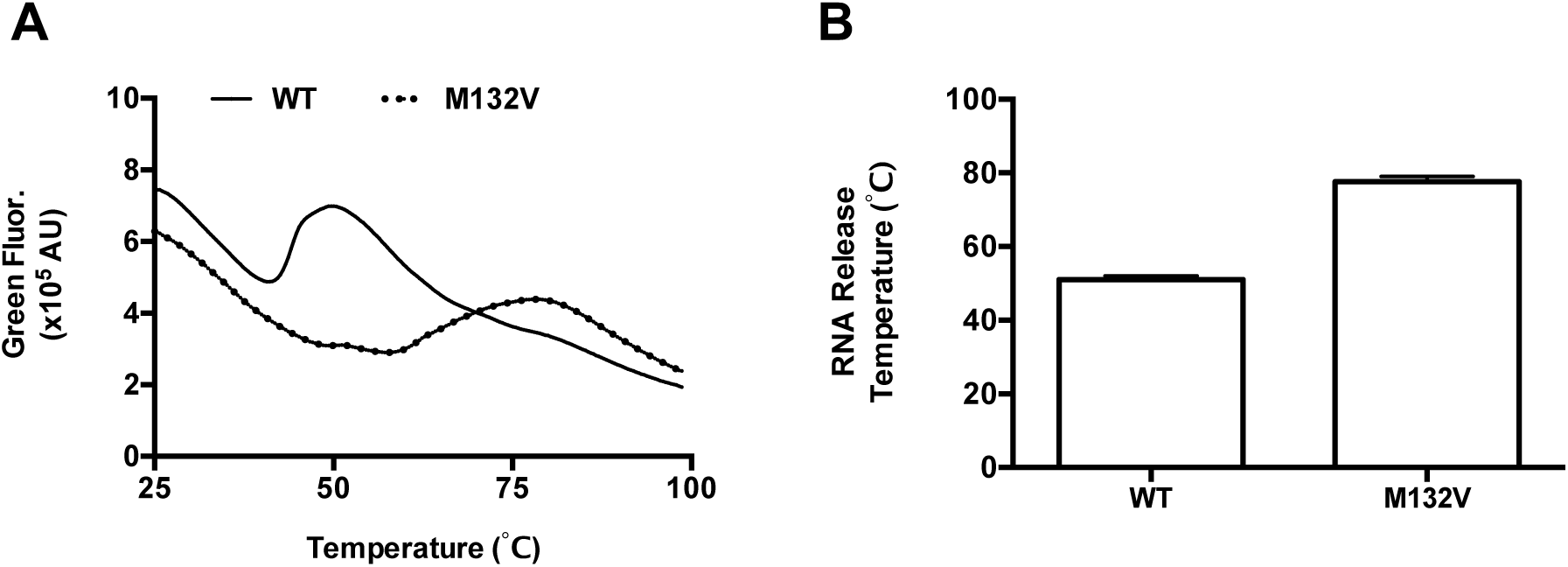
Particle Stability Thermo Release assay (PaSTRy). Gradient purified viruses were mixed with SYBR Green and placed in a real-time machine, where samples were heated from 25°C to 99°C on a 1% stepwise gradient with fluorescent monitoring. Peaks of SYBR Green fluorescence indicate virion RNA release. (A) Plots of SYBR Green fluorescence over varying temperatures. (B) Quantification of RNA release temperatures for three independent PaSTRy experiments. Data are mean ± SEM.

**FIG 3.**
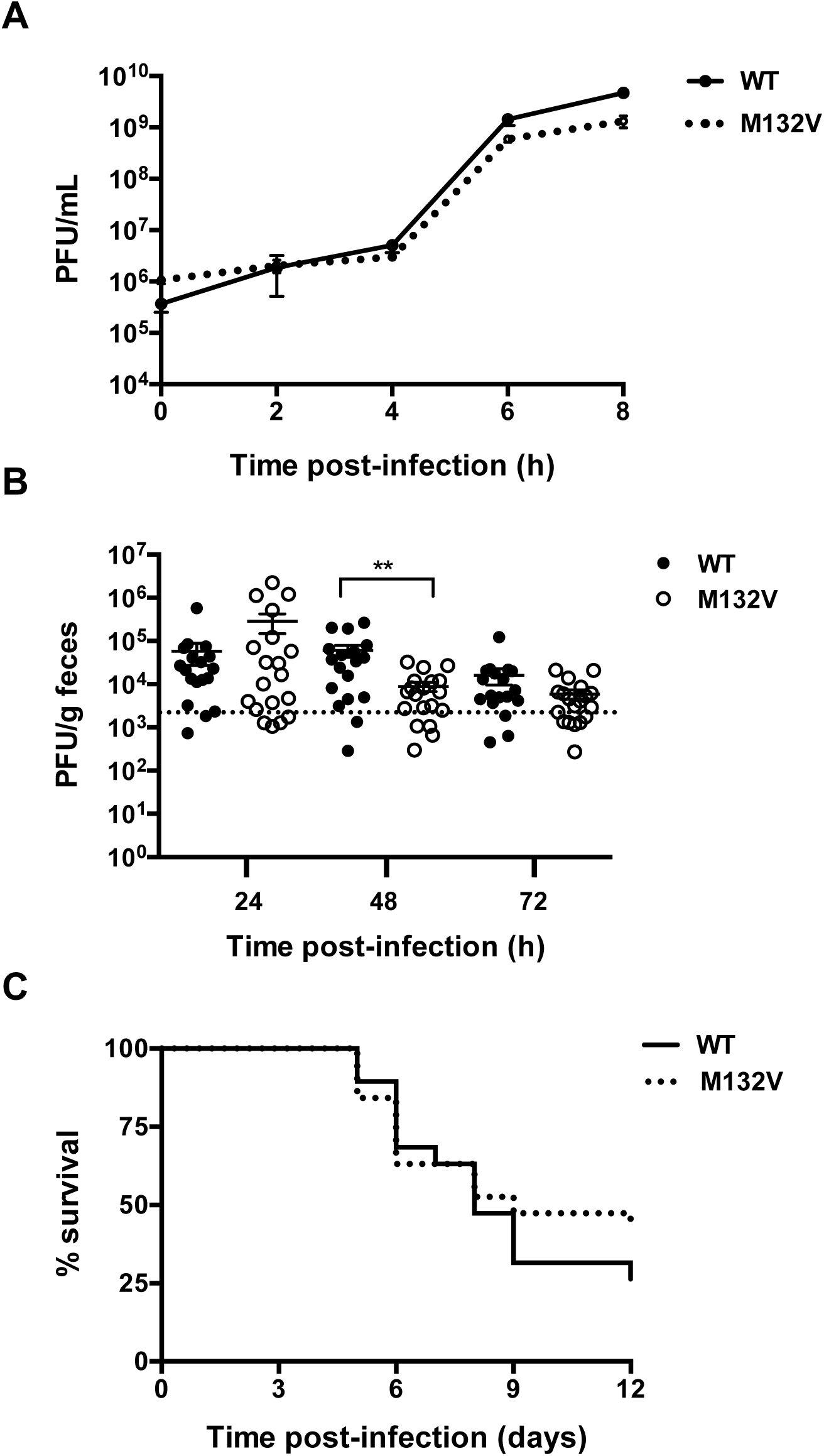
Comparison of WT and M132V PV replication and pathogenesis. (A) Single cycle growth curve assay. HeLa cells were infected with WT or M132V PV at an MOI of 10 at 37°C. Samples were harvested at various time points and yields were quantified by plaque assays; n=6. (B) Fecal shedding kinetics of WT and M132V viruses. C57BL/6 PVR-IFNAR^-/-^ mice were orally inoculated with 10^8^ PFU of WT or M132V viruses and feces were collected at 24, 48, and 72 hpi prior to quantification of viral titer by plaque assay. Dotted line represents the limit of detection. (C) Survival of C57BL/6 PVR-IFNAR^-/-^ mice following oral inoculation of WT or M132V poliovirus. Survival curves were not significantly different (p>0.5 by log rank test. Data in (B) and (C) are from two independent experiments with 18-20 animals for each virus. Data are mean ± SEM; **p<0.01.

**FIG 4.**
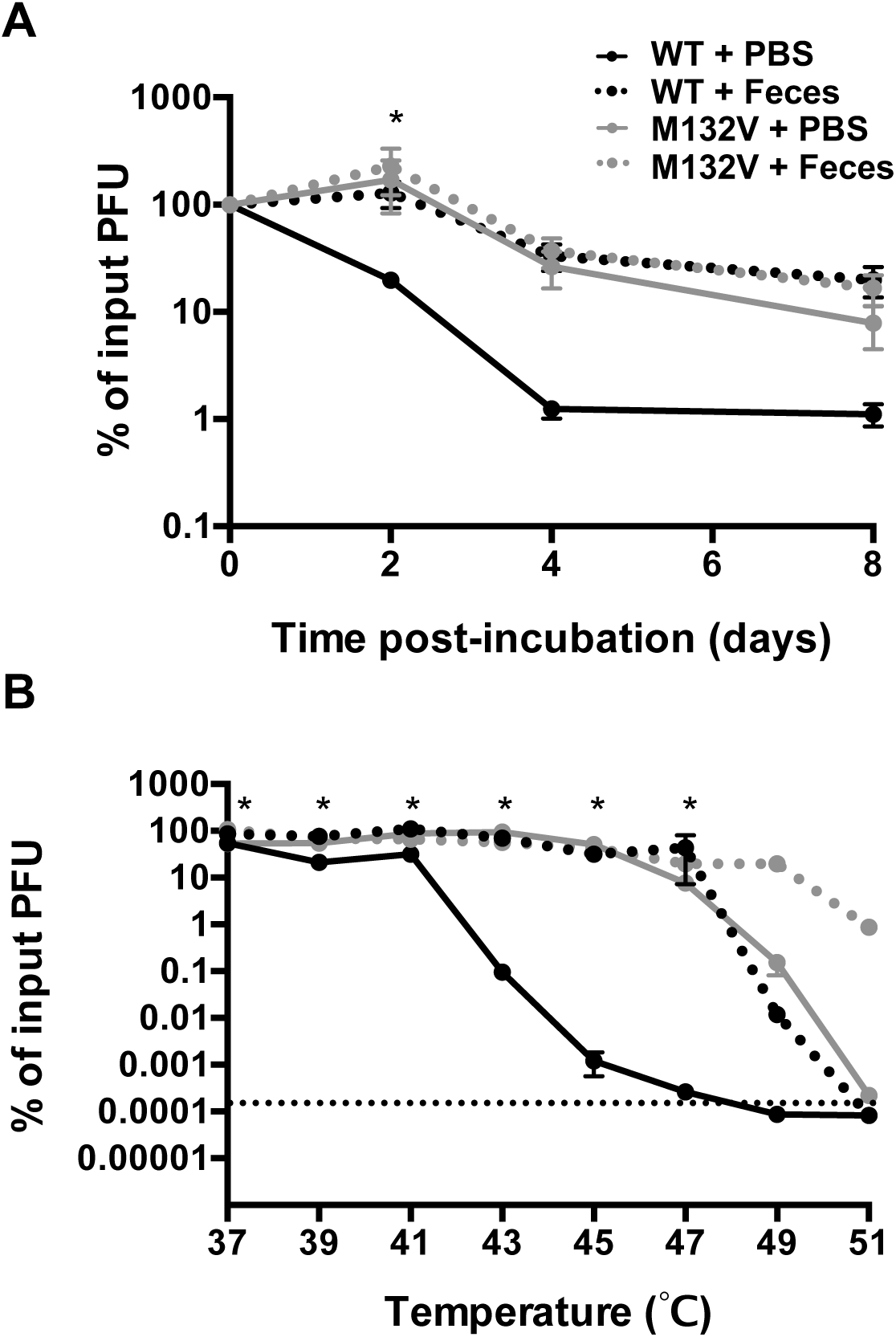
Thermal stability of WT and M132V PV in the presence or absence of feces. (A) Thermal stability profiles of WT and M132V PV at 37°C. Feces were collected from uninfected C57BL/6 PVR-IFNAR^-/-^ mice and resuspended in PBS prior to mixing with either WT or M132V PV. Samples were incubated at 37°C and plaque assays were performed on 0, 2, 4 and 8 d post-incubation, n=6. (B) Thermal stability profiles of WT and M132V PV at various temperatures in the presence or absence of feces during a 6 h incubation. Plaque assays were performed pre- and post-heat treatment, n=4. Data are mean ± SEM; Statistical analysis was performed using 2-way ANOVA. The * indicates that either one or of the more group is statistically significant (p<0.05) when compared to the WT PBS group. Data are mean ± SEM.

Since we selected for the M132V mutant at 43°C, it was possible that stability by fecal components was masked by the mutant’s inherent stability at 37°C. To investigate this, we examined the stability of WT and M132V PV during 6 h incubations at higher temperatures in the presence or absence of feces. As shown in Figure 4B, after 6 h incubation at 43°C in PBS, WT PV viability was reduced >1000-fold whereas M132V PV viability was reduced <2-fold. Feces stabilized WT PV up to an incubation temperature of 47°C. While M132V PV was stable in PBS up to 47°C, exposure to feces allowed recovery of 20% viable virus at 49°C and 1% viable virus at 51°C. Thus, for both WT and M132V viruses, feces stabilized viruses at temperatures that are damaging to the virions.

### WT and M132V PV have equivalent inactivation following bleach treatment

Hyper-stable virions could be dangerous to human health as they can persist longer in the environment and potentially resist inactivation by disinfectants such as bleach (12). Given that M132V PV is more thermostable than WT PV, we wanted to determine if the mutant is resistant to bleach inactivation. We treated 10^5^ PFU of either WT or M132V PV with 0.0001% bleach for 1 min, neutralized with sodium thiosulfate, and we performed plaque assays to determine the amount of viable virus before and after treatment. WT and M132V PV had equivalent inactivation following bleach treatment (Figure 5). This result suggests that the M132V mutation confers resistance to heat but not resistance to bleach, indicating that the mechanisms of virion inactivation by heat and bleach are distinct.

## DISCUSSION

Virion stability in the environment is crucial for transmission of PV and other enteric viruses. Previous work has shown that the microbiota can enhance PV replication, shedding, and pathogenesis in mice and that microbiota increase virion stability and cell attachment (12, 13). Specifically, our lab has shown that binding to bacterial surface components lipopolysaccharide or peptidoglycan enhanced PV thermostability. How bacteria promote virion stability is still unclear. To explore mechanisms and consequences of virion stabilization, we selected for and characterized a hyper-stable PV mutant.

**FIG 5.**
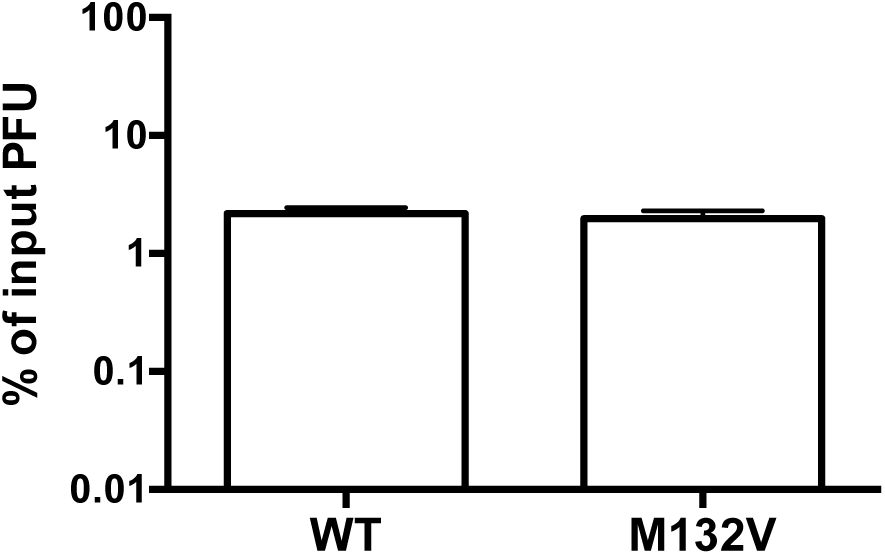
Stability of WT and M132V PV in the presence of bleach. WT or M132V PV were treated with 0.0001% fresh bleach for 1 min followed by neutralization. Plaque assays were performed to determine the amount of viable virus before and after treatment. Data are mean ± SEM.

Following repeated exposure to heat, we identified a single amino acid change, M132V in capsid protein VP1, that limited thermal activation of PV. VP1-M132 is buried in the capsid interior. Although M132 is located in a relatively conserved region of VP1, leucine is present in this position in other picornaviruses such as coxsackievirus B3, Aichi virus, and Mengo virus. In spite of the sequence conservation at M132 among PV isolates, M132V PV did not have major replication defects in culture cells or in mice (Figure 3).

Other thermostable PV mutants have been identified by other groups. For example, Adeyemi et al. and Shiomi et al. both found that VP1-V87A confers heat resistance and Adeyemi et al. also found that VP1-I194V confers heat resistance (8, 9). Ours is the first study to identify VP1-M132V as a stability determinant, and neither VP1-V87A or VP1-I194V were selected during our heat passage experiments. These results suggest that multiple amino acids in VP1 can contribute to heat resistance and that there are multiple paths to generate heat resistant PV variants.

Our work has shown that bacteria and bacterial glycans limit inactivation from both heat and bleach treatment, which led us to hypothesize that bacterial glycans bind virions and limit virion conformational changes and “breathing” that can lead to premature RNA release (12, 13). Although the specific binding site(s) of bacterial components on viral capsids have not been identified, it is highly likely that surface-exposed residues are involved. Importantly, M132V PV is not defective in bacterial binding (Aguilera et al., accompanying manuscript), and exposure to feces limits thermal inactivation of M132V PV at very high temperatures (49°C/51°C, Figure 4B). Thus, M132V PV does not require feces or bacteria for stabilization at a wide range of temperatures, but can be stabilized by feces at very high temperatures.

We originally sought virion stability mutants to gain insight into factors that affect capsid stability and to probe mechanisms and consequences of bacterial-mediated virion stabilization. Two lines of evidence suggest that mechanisms of stabilization by the M132V mutation and by bacteria/feces are different. First, if the two stabilization mechanisms were the same, the mutant should not be stabilized by bacteria or feces; however, the M132V mutant PV was stabilized by feces at very high temperatures (49°C/51°C, Figure 4B). This result suggests that virion thermal stabilization by bacteria is separate from and stronger than virion stabilization by the M132V mutation. Second, if the two stabilization mechanisms were the same, the M132V mutation should confer bleach resistance to the same extent as exposure to bacteria; however, the M132V mutant PV was as susceptible as WT PV to bleach inactivation (Figure 5). This result suggests that virion bleach stabilization by bacteria and bacterial glycans is separate from virion thermal stabilization by the M132V mutation. Overall our results suggest capsid dynamics are complex and that multiple distinct mechanisms influence PV stability and inactivation.

## EXPERIMENTAL METHODS

### Viruses and Cells

Serotype 1 Mahoney PV cell culture infections and plaque assays were performed using HeLa cells as previously described (17). HeLa cells were propagated in Dulbecco’s modified Eagle’s medium (DMEM) supplemented with 10% calf serum. Virus stocks from cellular lysates were derived from low passage stocks. For PaSTRy experiments, PV stocks were CsCl-gradient purified as previously described (13).

### Selection, sequencing, and cloning of M132V PV

To select for thermal stable PV mutants, 10^6^ PFU PV was incubated in PBS supplemented with 100 µg/ml CaCl_2_ and 100 µg/ml MgCl_2_ (PBS+) for 6 h at 43°C. Remaining viable viruses were amplified using HeLa cells 12-16 h (until cytopathic effects were visible. Sequential heating and amplification of viable viruses was performed a total of 10 times. Viral RNA was isolated using TRIzol as recommended by the manufacturer’s protocol (Sigma) and cDNA covering the capsid-coding region was synthesized using SuperScript II reverse transcriptase (Invitrogen). PCRs were performed to amplify the coding regions of VP1-4 and products were sequenced by the University of Texas Southwestern Sequencing Core. The sequenced region revealed one amino acid change, M132V (ATG to GTG) in VP1, and two silent mutations in VP1 in some isolates (T135: ACC to ACT and N203: AAC to AAT). To generate a virus containing only the VP1-M132V mutation, a 486-bp fragment containing the M132V mutation (and no other amino acid changes) was generated by SnaBI/NheI digestion of an RT-PCR product from the passage 10 virus. This fragment was cloned into a new PV plasmid at nucleotide 2470 (NheI) and 2956 (SnaBI). The new plasmid was sequenced and confirmed (nt 2470-2956). To generate M132V virus, the plasmid was transfected into HeLa cells along with a plasmid encoding the T7 DNA-dependent RNA polymerase to produce viral stocks as previously described (13).

### PV Thermal Stability Assays

Feces were collected from four-to six-week-old female C57BL/6 PVR-IFNAR-/-mice (16). Fecal pellets were resuspended in PBS+ to a final concentration of 0.0641g/mL. For titer-based stability assays in Figure 4A, 10^5^ PFU of PV was mixed with PBS+ or fecal slurry and incubated at 37°C for 2, 4, 6, or 8 days. For Figure 4B, 10^8^ PFU of PV was mixed with PBS+ or fecal slurry and incubated at the indicated temperatures for 6 h. Titers were quantified by plaque assays using HeLa cells. PV titers from pre- and post-heat treatment were calculated to determine the % input PFU. For PaSTRy experiments, one microgram of CsCl-gradient purified virus was mixed with SYBR GREEN II (10X final concentration) and buffer (10mM HEPES pH 8.0, 200mM NaCl) to a final volume of 30µl. Samples were heated from 25°C to 99°C on a 1% stepwise gradient with fluorescent monitoring using the ABI 7500 real-time instrument (12).

### Single-cycle growth curve assays

Six-well plates were seeded with 2x10^6^ HeLa cells/well. Cells were inoculated with either WT or M132V PV at a MOI of 10. After a 30 min incubation at 37°C, the inoculum was aspirated from the cells. Cell monolayers were washed with PBS and 3mL of fresh media was added. After 2, 4, 6, or 8 hpi at 37°C, cells were washed with PBS, trypsinized, and pelleted. Cells were freeze-thawed three times and the released intracellular viruses were quantified by plaque assay using HeLa cells.

### Mouse experiments

All animals were handled according to the Guide for the Care of Laboratory Animals of the National Institutes of Health. All mouse studies were performed at UT Southwestern (Animal Welfare Assurance #A3472-01) using protocols approved by the local Institutional Animal Care and Use Committee in a manner designed to minimize pain, and any animals that exhibited severe disease were euthanized immediately. Four-to five-week-old female and C57BL/6 PVR-IFNAR-/-mice were orally inoculated with 10^8.^PFU WT or M132V PV. Feces were collected at 24, 48, 72 hpi and viral shedding in feces was quantified by plaque assays. Survival of infected animals was also monitored for 12 dpi. Because PV disease is progressive and irreversible in this system, mice were euthanized at the first sign of disease, and these time points are shown in Fig. 3C.

### Bleach inactivation assay

Bleach inactivation assays were performed as previously described (12). Briefly, 10^5^ PFU of each virus was mixed with PBS+ and incubated at 37°C for 1 h. After incubation, samples were processed in 0.0001% fresh bleach (diluted immediately before the experiment) for 1 min. Bleach was neutralized by adding equal volume of 0.01% sodium thiosulfate (Sigma). Plaque assays were performed to determine amount of viable virus before and after treatment.

### Statistical analysis

The differences between groups were examined by unpaired two-tailed Student *t* tests. P< 0.05 was considered statically significant. WT and M132V mouse survival curves were not significantly different (p>0.5 by log rank test Figure 3C). The differences among the groups in the fecal stability assays (Figure 4) were assessed by two-way ANOVA tests.

## ACKNOWLEDGEMENTS

We thank Broc McCune and Arielle Woznica for critical review of the manuscript. We thank Nam Nguyen for assistance with structure models and Arielle Woznica for assistance with the animal studies.

## References

1. Stanway G. 1990. Structure, function and evolution of picornaviruses. J Gen Virol 71 (Pt 11):2483–2501.

2. Hogle JM. 2002. Poliovirus cell entry: common structural themes in viral cell entry pathways. Annu Rev Microbiol 56:677–702.

3. Rossmann MG, He Y, Kuhn RJ. 2002. Picornavirus-receptor interactions. Trends Microbiol 10:324–331.

4. Lin J, Lee LY, Roivainen M, Filman DJ, Hogle JM, Belnap DM. 2012. Structure of the Fab-labeled “breathing” state of native poliovirus. J Virol 86:5959–5962.

5. Li Q, Yafal AG, Lee YM, Hogle J, Chow M. 1994. Poliovirus neutralization by antibodies to internal epitopes of VP4 and VP1 results from reversible exposure of these sequences at physiological temperature. J Virol 68:3965–3970.

6. Roivainen M, Piirainen L, Rysä T, Närvänen A, Hovi T. 1993. An immunodominant N-terminal region of VP1 protein of poliovirion that is buried in crystal structure can be exposed in solution. Virology 195:762–765.

7. Ofori-Anyinam O, Vrijsen R, Kronenberger P, Boeyé A. 1995. Heat stabilized, infectious poliovirus. Vaccine 13:983–986.

8. Shiomi H, Urasawa T, Urasawa S, Kobayashi N, Abe S, Taniguchi K. 2004. Isolation and characterisation of poliovirus mutants resistant to heating at 50 degrees Celsius for 30 min. J Med Virol 74:484–491.

9. Adeyemi OO, Nicol C, Stonehouse NJ, Rowlands DJ. 2017. Increasing Type 1 Poliovirus Capsid Stability by Thermal Selection. J Virol 91.

10. De Palma AM, Vliegen I, De Clercq E, Neyts J. 2008. Selective inhibitors of picornavirus replication. Med Res Rev 28:823–884.

11. McSharry JJ, Caliguiri LA, Eggers HJ. 1979. Inhibition of uncoating of poliovirus by arildone, a new antiviral drug. Virology 97:307–315.

12. Robinson CM, Jesudhasan PR, Pfeiffer JK. 2014. Bacterial lipopolysaccharide binding enhances virion stability and promotes environmental fitness of an enteric virus. Cell Host Microbe 15:36–46.

13. Kuss SK, Best GT, Etheredge CA, Pruijssers AJ, Frierson JM, Hooper LV, Dermody TS, Pfeiffer JK. 2011. Intestinal microbiota promote enteric virus replication and systemic pathogenesis. Science 334:249–252.

14. Erickson AK, Jesudhasan PR, Mayer MJ, Narbad A, Winter SE, Pfeiffer JK. 2018. Bacteria Facilitate Enteric Virus Co-infection of Mammalian Cells and Promote Genetic Recombination. Cell Host Microbe 23:77–88.e5.

15. Walter TS, Ren J, Tuthill TJ, Rowlands DJ, Stuart DI, Fry EE. 2012. A plate-based high-throughput assay for virus stability and vaccine formulation. J Virol Methods 185:166–170.

16. Ida-Hosonuma M, Iwasaki T, Yoshikawa T, Nagata N, Sato Y, Sata T, Yoneyama M, Fujita T, Taya C, Yonekawa H, Koike S. 2005. The alpha/beta interferon response controls tissue tropism and pathogenicity of poliovirus. J Virol 79:4460–4469.

17. Kuss SK, Etheredge CA, Pfeiffer JK. 2008. Multiple host barriers restrict poliovirus trafficking in mice. PLoS Pathog 4:e1000082.

